# Accuracy of gene expression prediction from genotype data with PrediXcan varies across diverse populations

**DOI:** 10.1101/524728

**Authors:** Anna Mikhaylova, Timothy Thornton

## Abstract

Predicting gene expression with genetic data has garnered significant attention in recent years. PrediXcan is one of the most widely used gene-based association methods for testing imputed gene expression values with a phenotype due to the invaluable insight the method has shown into the relationship between complex traits and the component of gene expression that can be attributed to genetic variation. The prediction models for PrediXcan, however, were obtained using supervised machine learning methods and training data from the Depression and Gene Network (DGN) and the Genotype-Tissue Expression (GTEx) data, where the majority of subjects are of European descent. Many genetic studies, however, include samples from multi-ethnic populations, and in this paper we assess the accuracy of gene expression predictions with PrediXcan in diverse populations. Using transcriptomic data from the GEUVADIS (Genetic European Variation in Health and Disease) RNA sequencing project and whole genome sequencing data from the 1000 Genomes project, we evaluate and compare the predictive performance of PrediXcan in an African population (Yoruban) and four European populations. Prediction results are obtained using a range of models from PrediXcan weight databases, and Pearson’s correlation coefficient is used to measure prediction accuracy. We demonstrate that the predictive performance of PrediXcan varies across populations (F-test p-value < 0.001), where prediction accuracy is the worst in the Yoruban sample compared to European samples. Moreover, the performance of PrediXcan varies not only among distant populations, but also among closely related populations as well. We also find that the qualitative performance of PrediXcan for the populations considered is consistent across all weight databases used.

## 1 Introduction

In the past decade, genome-wide association studies (GWAS) have identified thousands of genetic variants significantly associated with a wide range of human phenotypes. The vast majority of these studies, however, were conducted in samples from European ancestry populations [1–5]. Differences in allele frequencies, genetic architecture, and linkage disequilibrium (LD) patterns across ancestries suggest that GWAS discoveries can fail to generalize across populations, and recent publications have provided compelling evidence that GWAS findings often do not transfer from European populations to other ethnic groups. For example, Carlson et al. analyzed multi-ethnic data from the PAGE Consortium and concluded that some GWAS-identified variants from European ancestry population had different magnitude and direction of allelic effects in non-European populations and the differential effects were more persistent in African Americans [6]. Moreover, genetic risk prediction models derived from European GWAS were unreliable when applied to other ethnic groups [6]. Martin et al. examined the impact of population history on polygenic risk scores and demonstrated that they were biased and confounded by population structure. [7]. Since genetic risk prediction accuracy depends on genetic similarity between the target and discovery cohorts, Martin et al. advised against interpreting the scores across populations and recommended computing them in genetically similar cohorts.

Associations between genetic variation and molecular traits, such as gene expression, have advanced our understanding of the mechanisms underlying trait-variant associations [8]. Prior studies have shown that a large proportion of GWAS variants identified for complex traits are expression quantitative trait loci (eQTLs): i.e., they play a role in regulating gene expression [9]. Thus, eQTLs can aid in prioritizing likely causal variants among the ones identified by GWAS, especially if they are found in non-coding regions, and uncover the mechanisms by which genotypes influence phenotypes [8]. So having three types of data - genotype, phenotype and gene expression - on the same set of subjects can be advantageous for investigating the relationships between phenotypes and genetic background of a subject and underlying processes. However, collecting all of these data types is often not feasible due to cost and tissue availability. Additionally, eQTL studies have the same pitfalls as GWASs - the majority of the detected eQTLs are not causal, but may be in LD with causal variants. Similar to variants identified through GWAS, eQTL findings might fail to replicate in diverse populations due to LD patterns that differ across populations.

Recently methods, such as PrediXcan, have been proposed for integrating eQTL studies and GWASs [10]. Such methods have multiple advantages over traditional GWAS methods, especially where expression data from the tissue of interest are not available and in cases when gene expression is in the causal pathway between genotypic variants and phenotype. PrediXcan can lead to an increase in power to detect associations for multiple reasons. First, it removes environmental noise and focuses on the genetically regulated component of gene expression. Second, PrediXcan bases gene expression prediction on a limited number of variants that are 1Mb upstream and downstream from the gene and then tests for association between the predicted expression and a phenotype. So, by including fewer variants that are potentially causal for every gene, the method has better power to detect eQTLs. Lastly, by conducting tests on aggregated variants instead of testing every variant, PrediXcan dramatically reduces multiple testing burden.

However, PrediXcan models were built using data from the Depression Genes and Networks (DGN) and the Genotype-Tissue Expression (GTEx) Project - both of which consist primarily of European-ancestry subjects. This poses the question of how accurate PrediXcan expression predictions are for non-European ancestry populations. Previous research has reported differences in gene expression levels across diverse populations from the HapMap3 project noting that 77% of eQTLs are population specific and only 23% are shared between two or more populations [11]. More distantly related populations have more differentially expressed genes, although this can often be explained by the expression of different gene transcripts across populations [12].

In this work, we investigated whether the predictive performance of PrediXcan differs across four European populations and one African populations using the Genetic European Variation in Health and Disease (GEUVADIS) [12] and 1000 Genomes Projects data [12,13]. We predicted gene expression levels using seven PrediXcan weight databases derived from whole blood and lymphoblastoid cell lines (LCL) expression data. To test prediction accuracy across populations, we compared observed and predicted gene expression levels by calculating Pearson’s correlation coefficients and then using linear mixed models to assess significant differences. In addition, we also evaluated the utility of whole-blood-based models when making predictions for LCL expression data. The results suggests that accuracy of PrediXcan for gene expression prediction differs across populations, even among closely related European ancestry populations. Furthermore, PrediXcan prediction accuracy is the worst in Africans across all weight databases we considered.

## 2 Materials and Methods

### 2.1 Datasets

We obtained gene expression data from the GEUVADIS Consortium and whole genome sequencing data from the 1000 Genomes Project. The gene expression data consisted of RNA sequencing on lymphoblastoid cell line (LCL) samples for 464 individuals from five populations. Of these, 445 subjects were in the 1000 Genomes Phase 3 dataset, including 358 subjects of European descent and 87 subjects of African descent. European samples included: Utah residents with Northern and Western European ancestry (CEU, *n* = 89), British individuals in England and Scotland (GBR, *n* = 86), Finnish in Finland (FIN, *n* = 92), and Toscani in Italy (TSI, *n* = 91). African samples included individuals of African descent from Yoruba in Ibadan, Nigeria (YRI, *n* = 87). Gene expression measurements were available for 23,722 genes.

We used seven PrediXcan weight databases: DGN whole-blood (further referred to as DGN), GTEx v6 1KG whole blood, GTEx v6 1KG LCL, GTEx v6 HapMap whole blood, GTEx v6 HapMap LCL, GTEx v7 HapMap whole blood (GTEx WB), and GTEx v7 HapMap LCL (GTEx LCL). The databases were downloaded from http://predictdb.org/.

### 2.2 Filtering out poorly predicted genes

Linear regression models were used to identify genes whose predicted values are not associated with the observed values at significance level of 0.05 in order to filter out the genes with poor prediction accuracy across all subjects. For each gene, we fit a linear regression model with observed gene expression as the outcome, and predicted gene expression as the predictor of interest. We performed the Wald test to assess the significance of the coefficient for each gene and excluded the genes whose corresponding p-values were above the significance level of 0.05.

We then calculated Pearson’s correlation coefficient, *r*, between observed and predicted expression values for every gene, in each population separately. A few genes had constant predicted gene expression levels across all subjects. Since we could not calculate the correlation if one of the variables was constant, we excluded those genes. Thus, for every gene we had five Pearson’s correlation coefficients, one per population. Note that we used *r* instead of the square of Pearson correlation, *r*^2^, in order to take the directionality of correlation into account. Using *r*^2^ as a measure of predictive accuracy can be misleading because a large proportion of genes predicted and observed expression values that are negatively correlated.

### 2.3 Prediction accuracy differences across populations and across tissues

To assess how the training of prediction models with different populations affects prediction accuracy, we used a linear mixed effect model approach. After filtering out poorly predicted genes, we fit the following model:

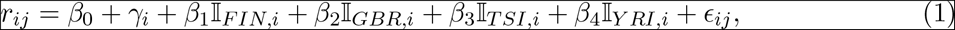

where *r*_ij_ is the correlation coefficient for gene *i* in population *j*; and 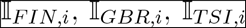, and 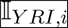 are indicator variables that are equal to 1 if the gene correlation was calculated on the population indicated in the subscript, and otherwise are equal to 0. Thus, we modeled population as a categorical predictor, with the CEU population as a reference. To account for variation between genes, we included a random intercept γ_*i*_ for each gene and we assumed that 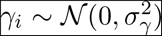. We also included an error term ϵ_*ij*_, such that 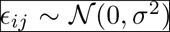. To simultaneously test for differences in correlation coefficients across populations, we used repeated measures ANOVA. To assess the association between the change in correlation coefficient and population, we tested the coefficients for each population using the likelihood-ratio test.

We also ran an additional analysis where we excluded the CEU population due to potentially lower quality of the CEU cell lines, as reported in the literature [14,15]. We fit a model identical to (1), excluding the CEU and using the FIN population as a reference:

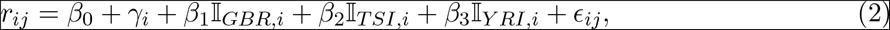

where the notation is the same as above. Again, we performed a repeated measures ANOVA to test for differences in gene correlations across the populations and the likelihood-ratio test to separately test the change in gene correlations for each population compared to the reference population.

To evaluate how PrediXcan performance with whole-blood (WB) databases differed from LCL databases, we restricted the set of genes to only those that were present in both the WB and LCL databases. We compared each pair of GTEx WB and GTEx LCL databases using a paired t-test. All the statistical analyses described above were performed in R version 3.3.3.

## 3 Results

### 3.1 Overview of PrediXcan weight databases

In Table 1, we summarize the main features of the PrediXcan weight databases that we used in the analyses. Compared to DGN database, GTEx databases have fewer gene models and smaller training sample sizes. HapMap and 1KG-based models differ in the number of variants used for training: GTEx Hapmap models were trained on the HapMap SNP set while GTEx 1KG were trained on the 1000 Genomes SNP set, so the latter utilize more SNPs when predicting expression. While GTEx LCL databases are based on relatively small training sets, they are derived from the same tissue as the GEUVADIS RNA-seq data we analyzed. Lastly, DGN and GTEx v7 sets of weights were trained only on the Europeans samples, while GTEx v6 databases had a small fraction of non-Europeans.

**Table 1:** Summary of PrediXcan databases used in analyses.

| PrediXcan Database | Training set size | Number of models | Number of SNPs used |
| --- | --- | --- | --- |
| DGN whole blood | 922 | 13,171 | 249,696 |
| GTEX v6 1KG whole blood | 338 | 6,759 | 185,786 |
| GTEX v6 1KG LCL | 114 | 3,759 | 125,045 |
| GTEX v6 HapMap whole blood | 338 | 6,588 | 136,941 |
| GTEX v6 HapMap LCL | 114 | 3,441 | 91,237 |
| GTEX v7 HapMap whole blood | 315 | 6,297 | 140,931 |
| GTEX v7 HapMap LCL | 96 | 3,045 | 88,143 |

To avoid repetition, we focus our attention on DGN, GTEx v7 WB and GTEx v7 LCL databases in the main text, and report our findings for the other four databases in the Supplementary material.

### 3.2 PrediXcan prediction accuracy differs across diverse populations

Using DGN, GTEx WB and GTEx LCL models and sequence data, we predicted gene expression for 10387, 5432 and 2777 genes, respectively (see Table 2). The number of genes with available predictions varied by population: the four European populations had similar counts and YRI had a slightly lower count. Because there was no variation in predicted expression values in at least one of the populations, we excluded 33 genes from DGN, 13 from GTEx WB, and 10 from GTEx LCL. From the remaining genes, we filtered out the ones with poor prediction accuracy based on associations between observed and predicted values, as described in the Materials and Methods section. Two-thirds of genes were excluded by this criteria from the genes predicted with DGN database, and slightly less than a half were excluded from gene sets predicted with the GTEx databases.

**Table 2:** Number of genes for which Pearson correlation coefficients are available by population and by PrediXcan weight database.

| PrediXcan database | DGN | GTEX v7 WB | GTEX v7 LCL |
| --- | --- | --- | --- |
| Genes with observed and predicted expression values | 10,387 | 5,432 | 2,777 |
| By population: |  |  |  |
| CEU | 10,385 | 5,432 | 2,777 |
| FIN | 10,385 | 5,432 | 2,777 |
| GBR | 10,385 | 5,432 | 2,777 |
| TSI | 10,385 | 5,432 | 2,776 |
| YRI | 10,354 | 5,419 | 2,767 |
| Genes before filtering | 10,354 | 5,419 | 2,767 |
| Genes after filtering | 3,493 | 2,288 | 1,699 |

Next, we computed gene correlation coefficients, separately in each of the five populations. Violin plots display the correlation coefficients by population across genes before and after filtering (see Figures 1A and 1B, respectively). We note that prediction accuracy is slightly lower for the African populations than for any of the European populations, regardless of the weight database used, and this trend is even more obvious after the filtering process.

**Figure 1:**
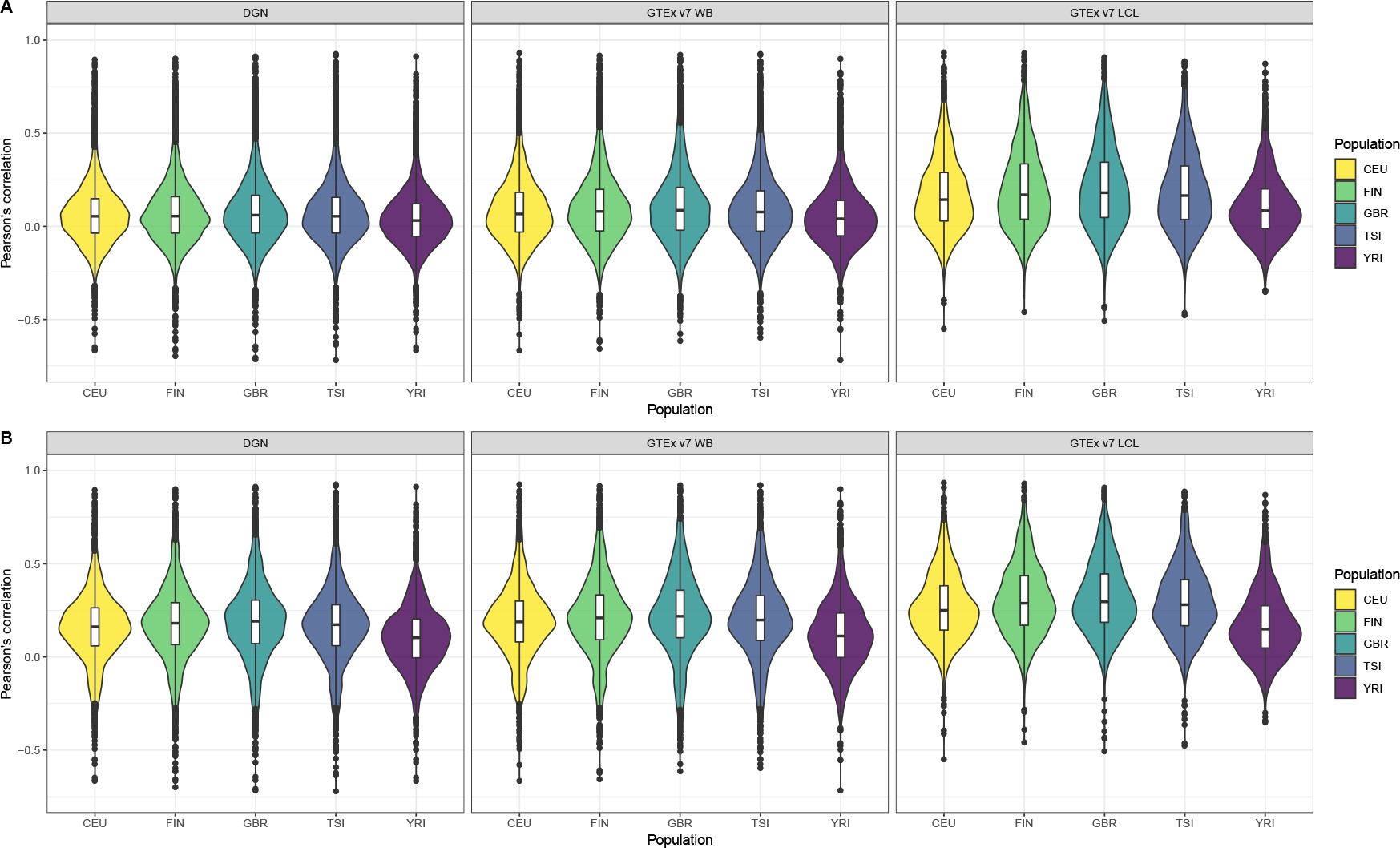
Violin plots of gene expression correlation coefficients by five populations using DGN, GTEx v7 WB and GTEx v7 LCL weight databases; **(A)** before and **(B)** after filtering out poorly predicted genes.

Afterwards, we binned the genes into six categories based on the gene correlation coefficients (see Table 3). The majority of genes have very poor prediction accuracy – of the genes predicted with whole-blood databases, a third have negative correlations and a half have correlations between 0 and 0.2. Of the genes predicted with LCL, a fifth have negative correlations and over a third have correlations between 0 and 0.2. The distribution of gene correlation coefficients is fairly similar across the four European populations, although predictive accuracy seems worse in CEU compared to FIN, GBR, and TSI. The predictive accuracy is the worst in the African sample. Across all populations, only a small number of genes were predicted with high accuracy (with *r* > 0.6). Furthermore, all European populations have a greater number of well-predicted genes than the African population, regardless of the weight database used.

**Table 3:** Binned gene correlation coefficients for the five populations using DGN, GTEx WB and GTEx LCL weight databases.

|  | Unfiltered |  |  |  |  | Filtered |  |  |  |  |
| --- | --- | --- | --- | --- | --- | --- | --- | --- | --- | --- |
|  | CEU | FIN | GBR | TSI | YRI | CEU | FIN | GBR | TSI | YRI |
| DGN database |  |  |  |  |  |  |  |  |  |  |
| $r < 0$ | 3,583 | 3,491 | 3,480 | 3,587 | 4,156 | 561 | 547 | 554 | 585 | 911 |
| $0 < r < 0.2$ | 5,107 | 4,976 | 4,812 | 4,954 | 5,001 | 1,533 | 1,379 | 1,258 | 1,409 | 1,674 |
| $0.2 < r < 0.4$ | 1,359 | 1,480 | 1,589 | 1,434 | 1,016 | 1,097 | 1,162 | 1,209 | 1,121 | 728 |
| $0.4 < r < 0.6$ | 239 | 302 | 354 | 290 | 147 | 236 | 300 | 353 | 289 | 146 |
| $0.6 < r < 0.8$ | 56 | 93 | 105 | 75 | 31 | 56 | 93 | 105 | 75 | 31 |
| $0.8 < r < 1$ | 10 | 12 | 14 | 14 | 3 | 10 | 12 | 14 | 14 | 3 |
| GTEx v7 WB database |  |  |  |  |  |  |  |  |  |  |
| $r < 0$ | 1,756 | 1,621 | 1,622 | 1,684 | 2,101 | 336 | 309 | 314 | 335 | 590 |
| $0 < r < 0.2$ | 2,471 | 2,450 | 2,366 | 2,456 | 2,491 | 877 | 786 | 732 | 820 | 993 |
| $0.2 < r < 0.4$ | 902 | 958 | 981 | 901 | 668 | 788 | 804 | 793 | 758 | 546 |
| $0.4 < r < 0.6$ | 210 | 282 | 329 | 278 | 117 | 207 | 281 | 328 | 275 | 117 |
| $0.6 < r < 0.8$ | 69 | 93 | 100 | 85 | 38 | 69 | 93 | 100 | 85 | 38 |
| $0.8 < r < 1$ | 11 | 15 | 21 | 15 | 4 | 11 | 15 | 21 | 15 | 4 |
| GTEx v7 LCL database |  |  |  |  |  |  |  |  |  |  |
| $r < 0$ | 546 | 488 | 484 | 509 | 774 | 80 | 69 | 55 | 69 | 274 |
| $0 < r < 0.2$ | 1,119 | 1,031 | 996 | 1,050 | 1,296 | 560 | 443 | 426 | 477 | 777 |
| $0.2 < r < 0.4$ | 718 | 742 | 761 | 736 | 510 | 675 | 681 | 692 | 681 | 461 |
| $0.4 < r < 0.6$ | 293 | 361 | 369 | 360 | 145 | 293 | 361 | 369 | 360 | 145 |
| $0.6 < r < 0.8$ | 80 | 126 | 137 | 96 | 38 | 80 | 126 | 137 | 96 | 38 |
| $0.8 < r < 1$ | 11 | 19 | 20 | 16 | 4 | 11 | 19 | 20 | 16 | 4 |

Next, we assessed the association between the prediction accuracy (as gene correlation coefficients) and population category via repeated measures ANOVA and linear mixed models. We present the parameter estimates and their 95% confidence intervals calculated using model-based standard errors for the model 1 in Table 4. Based on the repeated measures ANOVA, we find that prediction accuracy differs across populations, regardless of the weight database used (p-values for all databases were < 0.001). From the linear mixed model 1, we find that the prediction accuracy is significantly higher in FIN, GBR and TSI and significantly lower in YRI, compared to CEU (all p-values < 0.001). This suggests that predictive performance varies not only among distant populations, but also among closely related populations.

**Table 4:** Results from linear mixed models for population category (with CEU as a reference) and change in gene correlation coefficient among filtered genes.

|  | DGN |  |  | GTEx v7 WB |  |  | GTEx v7 LCL |  |  |
| --- | --- | --- | --- | --- | --- | --- | --- | --- | --- |
|  | Estimate | 95% CI | p-value | Estimate | 95% CI | p-value | Estimate | 95% CI | p-value |
| FIN | 0.019 | (0.014, 0.025) | < 0.001 | 0.021 | (0.015, 0.028) | < 0.001 | 0.038 | (0.030, 0.046) | < 0.001 |
| GBR | 0.029 | (0.023, 0.034) | < 0.001 | 0.032 | (0.025, 0.039) | < 0.001 | 0.051 | (0.043, 0.059) | < 0.001 |
| TSI | 0.010 | (0.004, 0.016) | < 0.001 | 0.013 | (0.007, 0.020) | < 0.001 | 0.027 | (0.019, 0.035) | < 0.001 |
| YRI | -0.054 | (-0.059, -0.048) | < 0.001 | -0.070 | (-0.077, -0.063) | < 0.001 | -0.097 | (-0.105, -0.089) | < 0.001 |

Finally, we repeated the analysis described above, this time excluding the CEU population. We present the parameter estimates and the corresponding 95% confidence intervals in Table 5. From the repeated measures ANOVA, we find that prediction accuracy differs across the four populations (p-values for all databases were < 0.001). Moreover, based on the coefficients and the corresponding p-values from the linear mixed model 2, we estimate the prediction accuracy to be significantly higher in GBR and significantly lower in TSI and YRI, compared to the FIN population (see corresponding p-values in Table 5). This difference in prediction accuracy is the greatest between YRI and FIN when GTEx v7 LCL weight database was used. Like in the analysis above, we notice that predictive performance differs across populations, including European populations.

**Table 5:**
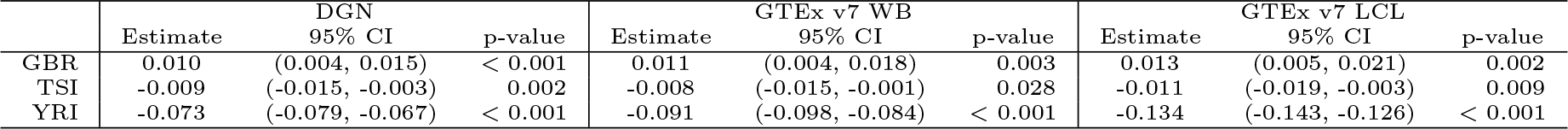
Results from linear mixed models for population category (excluding CEU, with FIN as a reference) and change in gene correlation coefficient among filtered genes.

|  | DGN |  |  | GTEx v7 WB |  |  | GTEx v7 LCL |  |  |
| --- | --- | --- | --- | --- | --- | --- | --- | --- | --- |
|  | Estimate | 95% CI | p-value | Estimate | 95% CI | p-value | Estimate | 95% CI | p-value |
| GBR | 0.010 | (0.004, 0.015) | < 0.001 | 0.011 | (0.004, 0.018) | 0.003 | 0.013 | (0.005, 0.021) | 0.002 |
| TSI | -0.009 | (-0.015, -0.003) | 0.002 | -0.008 | (-0.015, -0.001) | 0.028 | -0.011 | (-0.019, -0.003) | 0.009 |
| YRI | -0.073 | (-0.079, -0.067) | < 0.001 | -0.091 | (-0.098, -0.084) | < 0.001 | -0.134 | (-0.143, -0.126) | < 0.001 |

### 3.3 PrediXcan prediction accuracy differs between tissues

As can be seen in the violin plots in Figure 1, both databases based on whole blood perform similarly, and LCL-based database displays improved prediction accuracy. In order to compare pairwise gene correlations, we restricted our analyses to the 1,587 genes common in both GTEx v7 WB and GTEx v7 LCL. Scatter plots presented in Figure 2 suggest that the majority of genes have very similar correlation coefficients when using WB and LCL databases across all populations. However, we see more genes in the upper left corner, above the dotted line, indicating that using the LCL database results in more genes have better prediction accuracy. This result is not surprising since the expression data we used were derived from LCL. The results of the paired t-test are consistent with the visual examination of the data: the mean difference between gene correlations based on the GTEx v7 LCL model and based on the GTEx v7 WB model is 0.03 (p-value < 0.0001), with predictions based on the LCL model having better performance.

**Figure 2:**
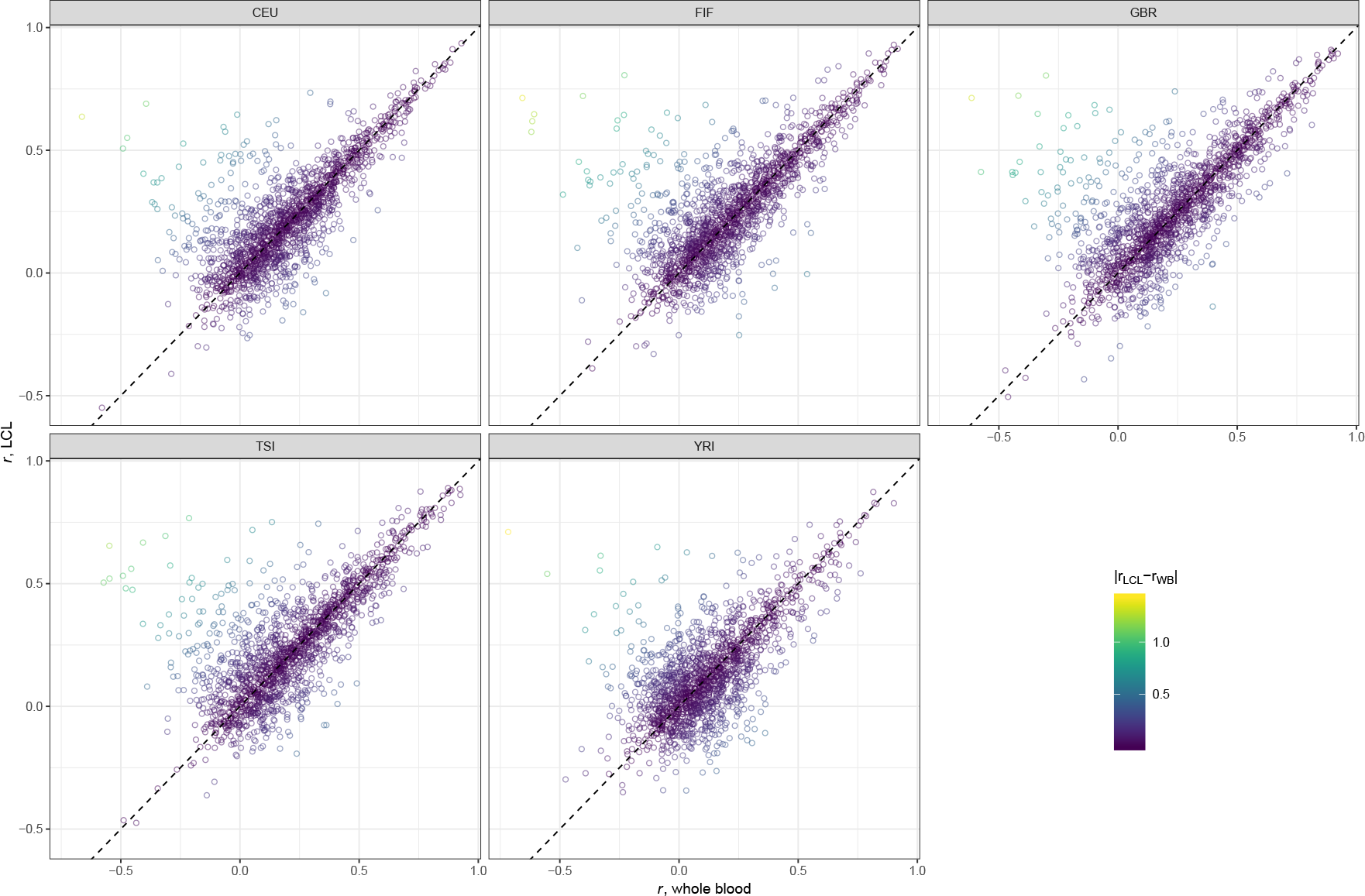
Scatter plots comparing gene correlation coefficients by population using GTEx v7 LCL vs GTEx v7 WB databases.

## 4 Discussion

In this work, we evaluated PrediXcan performance and compared it across five geographically diverse populations using multiple weight databases. Models from all seven weight databases were trained mostly on subjects of European ancestry; three of the databases were derived from LCL and the remaining four from whole blood. As a measure of prediction accuracy, we computed correlation coefficients for each gene in all populations and used the linear mixed models framework to quantify the differences in prediction performance across populations. We also investigated whether whole blood models could be used for predicting gene expression levels in LCL.

Overall, PrediXcan accurately predicted gene expression for some genes; however, the majority of genes had very poor correlation between measured and predicted expression levels. For almost half the genes, the correlation was negative. As expected, prediction accuracy was higher when the training and testing cohorts were of similar ancestry; i.e., models trained on Europeans performed better in the subjects of European descent and the worst in the African subjects. Surprisingly, prediction accuracy varied even among the European populations, with Finnish, British, and Italian populations having significantly higher accuracy than the CEU. These results held under all the weight databases we considered. Lastly, LCL-trained models outperformed whole-blood-trained models, although the prediction accuracy was similar for many of the genes.

A recent study reported consistent results to our findings and suggested that gene expression models should be trained on genetically similar populations [16]. Lack of genomic data from diverse populations limits the ability to effectively interpret and translate genomic results into clinical applications for individuals from admixed and other non-European populations. Our results in this paper emphasize the need to develop methods that account for ancestry and incorporate ancestral LD structure and allele frequencies differences. We also corroborate the importance of including more ancestrally diverse individuals in medical genomics to ensure that everyone gets the benefits of precision medicine and to avoid further exacerbating healthcare inequality.

We conclude this paper with some important caveats. LCLs are derived from B cells found in whole blood, and they provide a continuous supply of genetic material for GWAS and gene expression studies. However, they do undergo a transformation to become immortal that can change their biology and they do not have the same properties as native tissue [17]. Storage conditions, freeze-thaw cycles, and maturity of cell lines can also affect gene expression patterns [14,15]. The CEU cell lines were collected much earlier than the other cell lines and LCL age can have a confounding effect and bias downstream analyses [14]. This factor could have contributed to the differences in prediction accuracy among European populations. Lastly, our study had modest sample sizes and only one non-European population. Future work is needed to investigate the performance and prediction accuracy of PrediXcan and other related approaches for gene expression in other multi-ethnic and ancestrally diverse populations.

## Author Contributions

AM and TT conceived the idea, designed the analysis, interpreted the results, and wrote the paper. AM ran the analysis.

## Data Availability Statement

GEUVADIS expression data is avalable at Array Express (E- MTAB-264 and E-GEUV-1) at https://www.ebi.ac.uk/arrayexpress/experiments/ and 1000 Genomes project genotype data is available at http://www.internationalgenome.org/.

