## SupplementaryMaterial for "Accuracy of gene expression prediction from genotype data with PrediXcan varies across diverse populations"

### 1 Supplementary Tables and Figures

We present the results for the four PrediXcan weight databases omitted from the main text. The databases are GTEx v6 1KG whole blood (GTEx v6 1KG WB), GTEx v6 1KG LCL (GTEx v6 1KG LCL), GTEx v6 HapMap whole blood (GTEx v6 HM WB), and GTEx v6 HapMap LCL (GTEx v6 HM LCL).

Table 1: Number of genes for which Pearson correlation coefficients are available by population and by GTEx v6 database.

| PrediXcan database | 1KG WB | 1KG LCL | HM WB | HM LCL |
|---|---|---|---|---|
| Genes with observed and predicted expression values | 6,179 | 3,662 | 6,039 | 3,363 |
| By population: | | | | |
| CEU | 6,136 | 3,636 | 6,017 | 3,361 |
| FIN | 6,132 | 3,637 | 6,020 | 3,361 |
| GBR | 6,133 | 3,637 | 6,018 | 3,361 |
| TSI | 6,141 | 3,637 | 6,025 | 3,361 |
| YRI | 6,039 | 3,612 | 5,997 | 3,351 |
| Genes before filtering | 6,010 | 3,604 | 5,978 | 3,350 |
| Genes after filtering | 2,198 | 1,889 | 2,207 | 1,847 |

Table 2: Binned gene correlation coefficients for the five populations using GTEx v6 weight databases.

| | Unfiltered | | | | | Filtered | | | | |
|---|---|---|---|---|---|---|---|---|---|---|
| | CEU | FIN | GBR | TSI | YRI | CEU | FIN | GBR | TSI | YRI |
| | GTEx v6 1KG WB database | | | | | | | | | |
| $r < 0$ | 2,094 | 2,031 | 1,993 | 2,051 | 2,285 | 347 | 313 | 329 | 334 | 492 |
| $0 < r < 0.2$ | 2,804 | 2,747 | 2,673 | 2,770 | 2,826 | 870 | 809 | 730 | 804 | 976 |
| $0.2 < r < 0.4$ | 856 | 892 | 961 | 867 | 708 | 727 | 737 | 757 | 739 | 540 |
| $0.4 < r < 0.6$ | 183 | 249 | 280 | 240 | 147 | 181 | 248 | 279 | 239 | 146 |
| $0.6 < r < 0.8$ | 64 | 82 | 88 | 69 | 39 | 64 | 82 | 88 | 69 | 39 |
| $0.8 < r < 1$ | 9 | 9 | 15 | 13 | 5 | 9 | 9 | 15 | 13 | 5 |
| | GTEx v6 1KG LCL database | | | | | | | | | |
| $r < 0$ | 841 | 806 | 799 | 804 | 1030 | 97 | 99 | 86 | 100 | 236 |
| $0 < r < 0.2$ | 1,570 | 1,492 | 1,460 | 1,544 | 1,666 | 673 | 570 | 551 | 612 | 817 |
| $0.2 < r < 0.4$ | 849 | 841 | 833 | 829 | 676 | 775 | 755 | 740 | 750 | 604 |
| $0.4 < r < 0.6$ | 267 | 339 | 376 | 320 | 178 | 267 | 339 | 376 | 320 | 178 |
| $0.6 < r < 0.8$ | 69 | 112 | 117 | 89 | 48 | 69 | 112 | 117 | 89 | 48 |
| $0.8 < r < 1$ | 8 | 14 | 19 | 18 | 6 | 8 | 14 | 19 | 18 | 6 |
| | GTEx v6 HapMap WB database | | | | | | | | | |
| $r < 0$ | 2,092 | 2,052 | 1,986 | 2,065 | 2,246 | 350 | 331 | 329 | 368 | 504 |
| $0 < r < 0.2$ | 2,770 | 2,705 | 2,679 | 2,760 | 2,883 | 885 | 804 | 763 | 830 | 1,004 |
| $0.2 < r < 0.4$ | 879 | 892 | 942 | 846 | 685 | 736 | 743 | 745 | 703 | 535 |
| $0.4 < r < 0.6$ | 176 | 248 | 279 | 229 | 128 | 175 | 248 | 278 | 228 | 128 |
| $0.6 < r < 0.8$ | 55 | 72 | 79 | 68 | 33 | 55 | 72 | 79 | 68 | 33 |
| $0.8 < r < 1$ | 6 | 9 | 13 | 10 | 3 | 6 | 9 | 13 | 10 | 3 |
| | GTEx v6 HapMap LCL database | | | | | | | | | |
| $r < 0$ | 750 | 691 | 676 | 688 | 947 | 87 | 77 | 89 | 81 | 256 |
| $0 < r < 0.2$ | 1,471 | 1,405 | 1,365 | 1,434 | 1,568 | 689 | 593 | 536 | 601 | 820 |
| $0.2 < r < 0.4$ | 810 | 797 | 847 | 834 | 628 | 752 | 720 | 760 | 771 | 564 |
| $0.4 < r < 0.6$ | 257 | 350 | 328 | 299 | 164 | 257 | 350 | 328 | 299 | 164 |
| $0.6 < r < 0.8$ | 53 | 93 | 119 | 78 | 39 | 53 | 93 | 119 | 78 | 39 |
| $0.8 < r < 1$ | 9 | 14 | 15 | 17 | 4 | 9 | 14 | 15 | 17 | 4 |

Table 3: Results from linear mixed models for population category (with CEU as a reference) and change in gene correlation coefficient among filtered genes.

| Regression | GTEx v6 1KG WB | | GTEx v6 1KG LCL | | GTEx v6 HM WB | | GTEx v6 HM LCL | |
|---|---|---|---|---|---|---|---|---|
| parameter | Estimate | 95% CI | Estimate | 95% CI | Estimate | 95% CI | | |
| FIN | 0.025 | (0.018, 0.032) | 0.031 | (0.024, 0.039) | 0.020 | (0.013, 0.027) | 0.034 | (0.026, 0.041) |
| GBR | 0.031 | (0.023, 0.038) | 0.043 | (0.035, 0.050) | 0.028 | (0.021, 0.035) | 0.044 | (0.036, 0.051) |
| TSI | 0.015 | (0.008, 0.022) | 0.023 | (0.016, 0.031) | 0.011 | (0.004, 0.018) | 0.027 | (0.019, 0.034) |
| YRI | -0.040 | (-0.047, -0.033) | -0.061 | (-0.069, -0.054) | -0.047 | (-0.054, -0.040) | -0.064 | (-0.071, -0.056) |

Table 4: Results from linear mixed models for population category (excluding CEU, with FIN as a reference) and change in gene correlation coefficient among filtered genes.

| Regression | GTEx v6 1KG WB | | GTEx v6 1KG LCL | | GTEx v6 HM WB | | GTEx v6 HM LCL | |
|---|---|---|---|---|---|---|---|---|
| parameter | Estimate | 95% CI | Estimate | 95% CI | Estimate | 95% CI | | |
| GBR | 0.006 | (-0.001, 0.013) | 0.012 | (0.004, 0.019) | 0.008 | (0, 0.015) | 0.010 | (0.002, 0.018) |
| TSI | -0.009 | (-0.017, -0.002) | -0.008 | (-0.016, 0) | -0.010 | (-0.017, -0.002) | -0.007 | (-0.015, 0) |
| YRI | -0.064 | (-0.072, -0.057) | -0.092 | (-0.100, -0.085) | -0.067 | (-0.075, -0.060) | -0.099 | (-0.107, -0.091) |

### 1.1 Figures

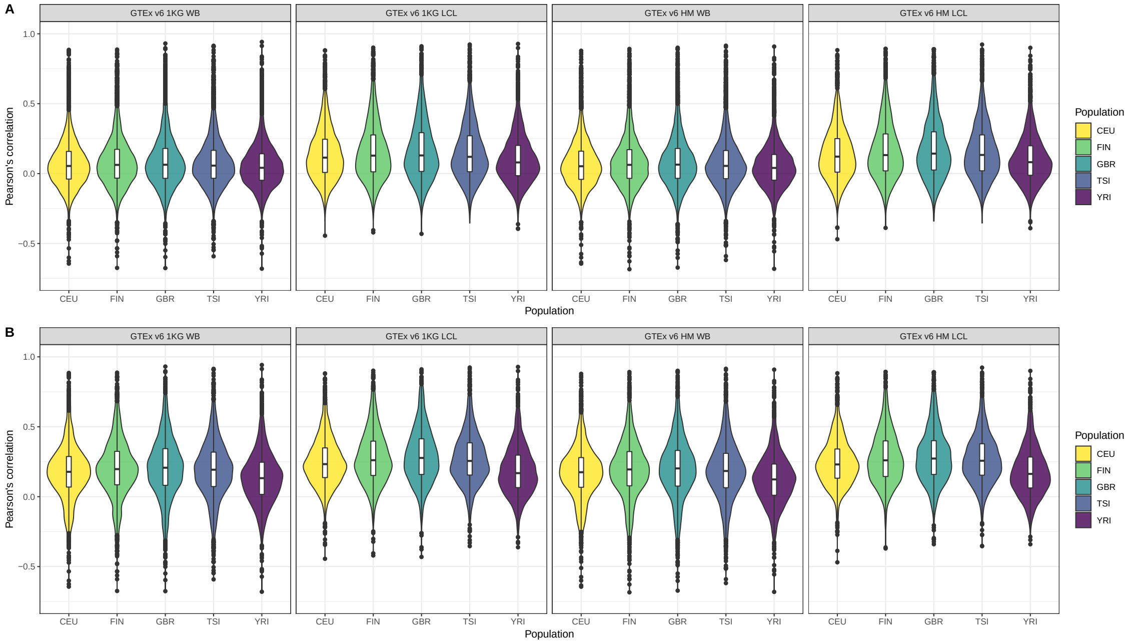


Figure 1: Violin plots of gene expression correlation coefficients by five populations using GTEx weight databases; **(A)** before and **(B)** after filtering out poorly predicted genes.

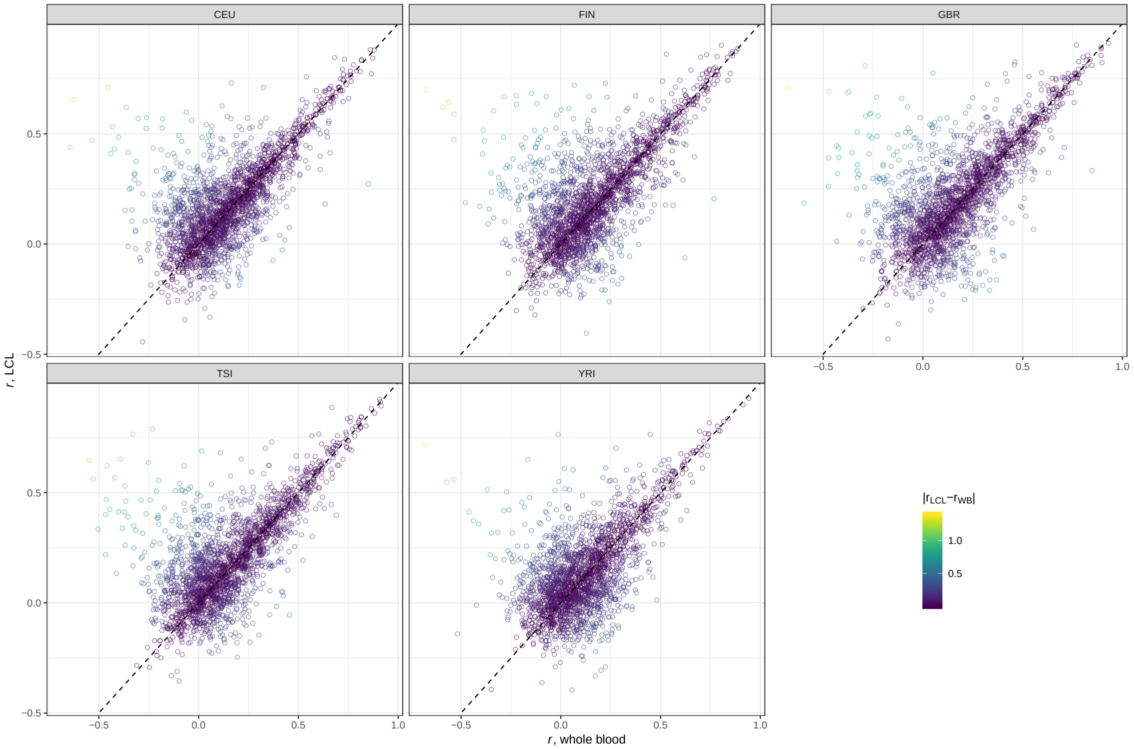


Figure 2: Scatter plots comparing gene correlation coefficients by population using GTEx v6 1KG LCL vs GTEx v6 1KG WB databases.

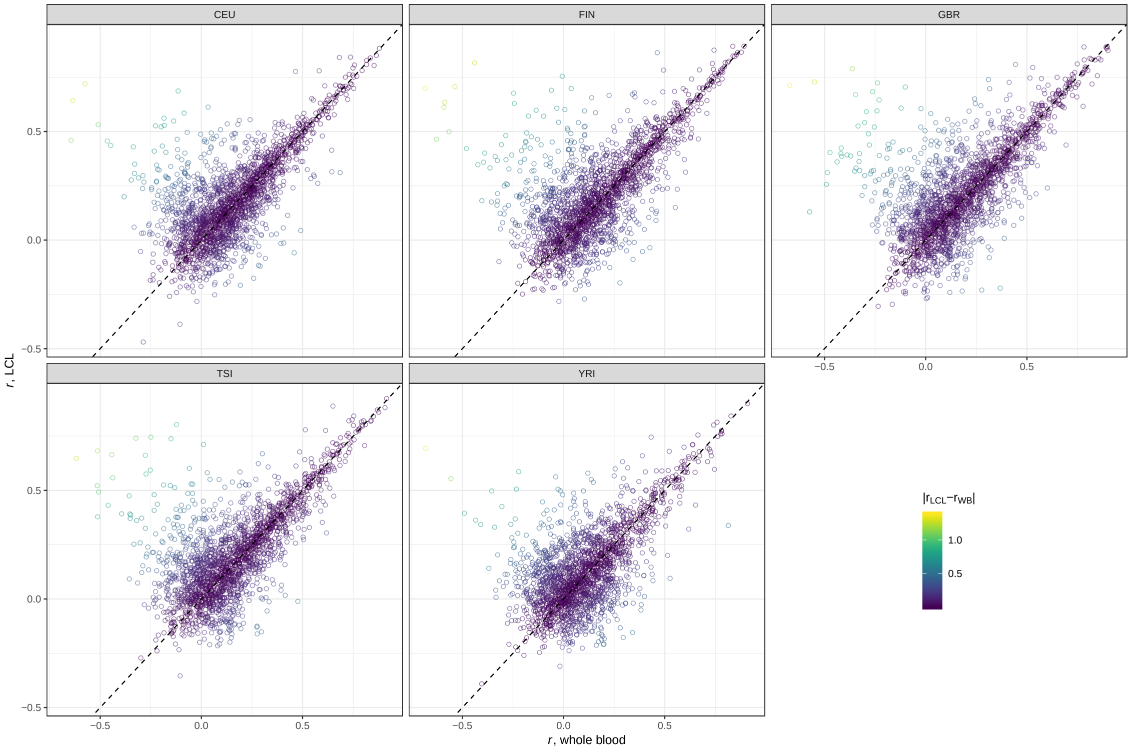


Figure 3: Scatter plots comparing gene correlation coefficients by population using GTEx v6 HapMap LCL vs GTEx v6 HapMap WB databases.
